# High-content interrogation of human induced pluripotent stem cell-derived cortical organoid platforms

**DOI:** 10.1101/697623

**Authors:** Madel Durens, Jonathan Nestor, Kevin Herold, Robert F. Niescier, Jason W. Lunden, Andre W. Phillips, Yu-Chih Lin, Michael W. Nestor

## Abstract

The need for scalable and high-throughput approaches to screening using 3D human stem cell models remains a central challenge in using stem cell disease models for drug discovery. It is imperative to develop standardized systems for phenotypic screening, yet most researchers screen cells across different platforms using a multitude of assays. In this study, we have developed a workflow centered on a small array of assays that can be employed to screen 3D stem cell cultures across a set of platforms. This workflow can be used as a starting point for a standardized approach to phenotypic screening. In this manuscript we hope to provide a roadmap for groups looking to start high-content screening using 3D organoid systems. To do this, we employ serum-free embryoid bodies (SFEBs) created from human induced pluripotent stem cells (hiPSCs). SFEBs are used in this study because they do not display the same level of heterogeneity observed in other neural organoid systems and they are amenable to high content imaging without cryosectioning. They contain populations of excitatory and inhibitory neurons that form synaptically active networks^1^ and medium- to high-throughput electrophysiology can be performed using SFEBs via the multielectrode array (MEA). The assays outlined in this study allow SFEBs to be scanned for neurite outgrowth, cell number and electrophysiological activity. SFEBs derived from control and disease hiPSCs can be used in combination with high-throughput screening assays to generate sufficient statistical power to compensate for the biological and experimental variability common in 3D cultures, while significantly decreasing processing speed, thus making this an efficient starting point for phenotypic drug screening.

## Introduction

Three-dimensional (3D) cell culture models derived from hiPSCs provide a promising avenue for studying disease mechanisms and for drug discovery because they can potentially recapitulate aspects of *in vivo* physiology more accurately than 2D systems. Previous studies have shown that 3D and 2D cultures vary greatly in terms of cell proliferation, differentiation, migration, cytoarchitecture, oxygen and nutrient gradients, and cell-cell and cell-extracellular matrix interactions^2,3^. These altered conditions impact cellular phenotypes and drug responses, and could ultimately lead to high failure rates in early stages of phenotypic drug screens performed solely in 2D cultures^4^.

More recently, the use of hiPSC to generate neural 3D cultures has improved our ability to model neurodevelopmental disorders with complex genetic and environmental etiologies, striking a balance between the relative simplicity of 2D cultures and the limited genetic heterogeneity and immense cost of animal models^5^. Brain organoids self-organize into neuroepithelial structures analogous to early fetal development, and form electrically-active networks. Several protocols have been developed to convey regional identities by the addition of patterning factors. Studies also have shown that neurons in 3D cultures are more electrically robust, and express more genes involved in neurological processes^5–7^. Many different methods for generating 3D cultures have emerged, including spheroid cultures, spinning bioreactors, microfluidic systems, and biopolymer scaffolds^2^. However, the utility of these culture methods, particularly in drug screening pipelines, has been hampered by a high degree of experimental and biological heterogeneity, increased cost, and the lack of validated methodologies for quantitative analysis^8,9^. 3D systems that model the development of cortical networks are typically difficult to work with in imaging and electrophysiological applications at scale because the tissue thickness requires cryosectioning and other special handling^7^. The development and adoption of medium-to high-throughput strategies for assaying 3D culture models is paramount to overcoming these challenges.

A technique developed by Nestor et al^1^ creates 3D SFEBs that approximate cortical development while also facilitating imaging and electrophysiological recording without the need for further processing. By transferring SFEBs onto cell-culture inserts composed of the polytetrafluoroethylene (PTFE) after aggregation, SFEBs thin to ~100 μm allowing high-throughput imaging in multi-well plates while maintaining synaptic connections^1^. SFEBs contain a heterogeneous population of calretinin- and calbindin-positive interneurons, vesicular glutamate transporter (VGlut) positive excitatory neurons and Tbr1-positive glutamatergic neural progenitors, and astrocytes, and can be used to model early cortical networks^1,10,11^. SFEBs also provide a consistent platform for working with 3D systems because they are less variable than other cerebral organoids^11^. The SFEBs used in this method are derived from peripheral blood mononuclear cells (PBMCs) and retain the original genetics and disease susceptibility of the individual from which they were derived; thus they are an excellent resource for *in vitro* disease modelling^12^. SFEBs derived from hiPSCs from control and disease cohorts can be used in combination with high-throughput screening assays to generate sufficient statistical power to compensate for the high variability observed in cerebral organoid systems ^8^.

Here we developed a framework and protocol to take advantage of the benefits of 3D systems, while compensating for their deficiencies. High-content imaging was utilized to analyze SFEBs for neurite outgrowth and morphology, cell population, and multiple other parameters. As SFEBs are electrophysiologically active, electrophysiological measurements were obtained in parallel via MEA to measure spontaneous and evoked spiking as well as long-term potentiation to model cortical network aberrations. In addition, Ca^2+^ imaging can be used as an analogue for single-cell action potentials^13^. These measurements can be taken before and after pharmacological manipulation allowing for the assaying of drug effects and to examine receptor function and synaptic plasticity. We describe a novel workflow to allow interrogation of SFEBs using three multiplex high-throughput techniques, namely immunohistochemistry, multi-electrode array (MEA) for electrophysiology, and Ca^2+^ imaging, on ArrayScan high-throughput technology (Figure 1).

**Figure 1.**
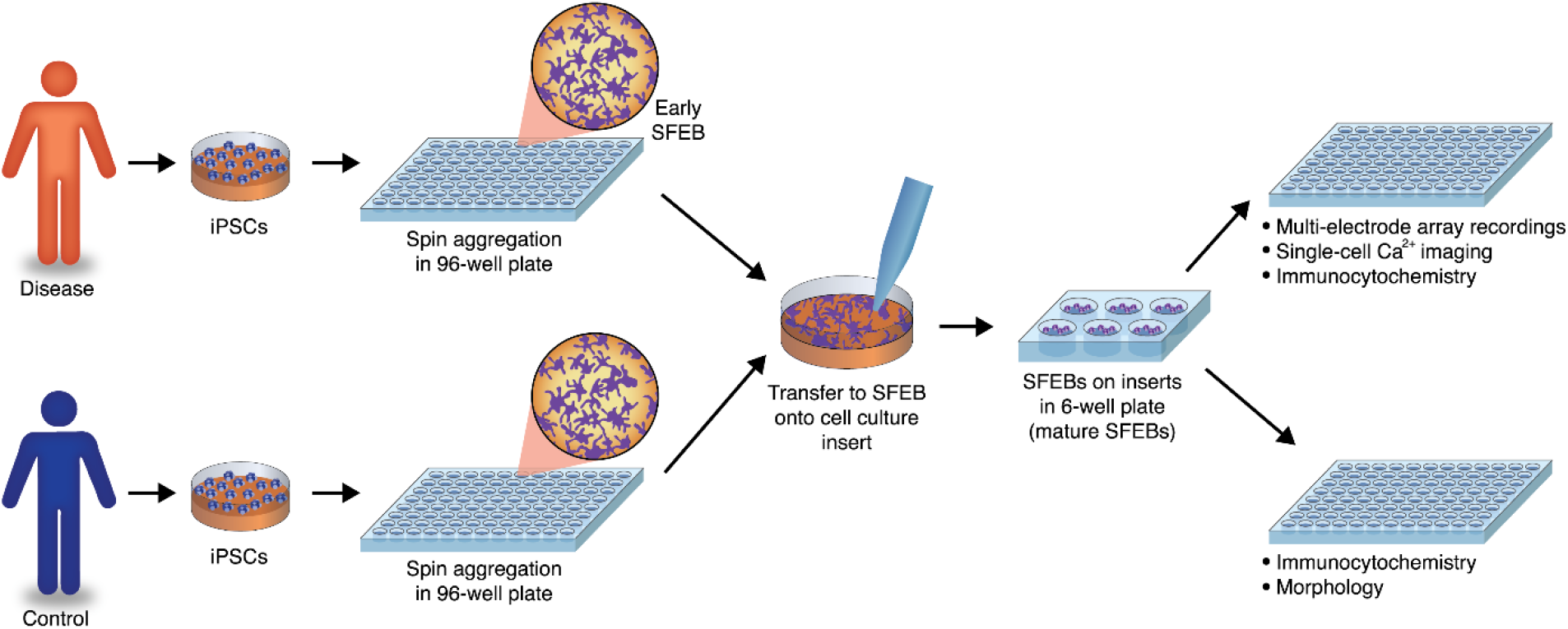
General protocol for generating SFEBs and using them for cell-based assays. The hiPSCs were derived from peripheral blood mononuclear cells (PBMCs, i.e. lymphocytes and monocytes) using previously described protocols.^14^ SFEBs were formed by centrifuging 20,000 cells per well on 96-well V-bottom plates, and allowing them to grow for 14 days. Early SFEBs were transferred to cell culture inserts within 6-well plates on day 15 and remained there through day 54.

## Materials and Methods

### Generation of SFEBs from hiPSCs

The hiPSCs were derived from PBMCs and maintained on a gamma-irradiated mouse embryonic fibroblast (MEF, ThermoFisher) feeder layer in mTeSR (StemCell Technologies) media supplemented with FGF (40 ng/ml, StemGent), CHIR99021 (10 μM, Stemgent), Thiazovivin (1μM, Stemgent), and PD0325901 (1 μM, Stemgent)^14^. The method used for generating SFEBs was adapted from Nestor et al^10^. Undifferentiated hiPSC colonies were manually picked and dissociated with Accutase (ThermoFisher). Dissociated cells were resuspended in media supplemented with Rho Khinase inhibitor (10 μM, StemGent) and plated for 1 hour in 10 cm-culture dishes coated with 0.1% gelatin solution (Millipore Sigma) to remove MEFs. The supernatant was subsequently plated in a 96-well V-bottom at a density of 20,000 cells with Neural Induction Medium (StemCell Technologies) supplemented with dual SMAD inhibitors SB321542 (10 μM, StemGent) and Dorsomorphin (1 μM, StemGent), 1 μM Thiazovivin, and 40 ng/ml FGF. Plates were then centrifuged at 163 x g for 3 minutes to promote aggregation.

After 14 days in vitro (DIV14), aggregates were transferred to 0.4μm PTFE cell culture inserts (Millipore) with Neural Formation medium (1:1 DMEM/F12, Neurobasal, 2% B27 supplement, 0.5% N2 supplement, 0.5% Non-essential amino acids, 1% SATO mix, 1% Insulin Transferrin Selenium A, 1% Pen/Strep) supplemented with neurotrophic factors (20 ng/ml BDNF (Peprotech), 20 ng/ml NT3 (Peprotech), 20 ng/ml β-NGF (Peprotech), 20 ng/ml FGF (StemGent), 1 μg/ml laminin (ThermoFisher), 5 μM forskolin (Sigma), 2 μg/ml heparin (Sigma)). Each insert was used for 8-12 SFEBs to allow adequate space for growth. At DIV30, terminal differentiation was induced by addition of 2 μM DAPT (StemGent), and withdrawal of FGF, β-NGF and heparin. DAPT was withdrawn from the media at DIV45. Media was changed 50% every other day until assays were performed.

### Neuronal Morphology

SFEBs at DIV60 were fixed using 4% paraformaldehyde for 30 minutes and transferred to optically clear 96 well U-bottom plates (Corning) to facilitate imaging. SFEBs were washed 3x with PBS and incubated with blocking solution (PBS with 5% Normal Donkey Serum (Jackson Immunoresearch Labs) and 0.1% Triton-X) for one hour. After blocking solution was removed, SFEBs were incubated with rabbit anti-VGlut primary antibody (1:250, Synaptic Systems) in blocking solution overnight at 4°C, then washed 3x with PBS with 0.1% Triton-X. This was followed by secondary antibody incubation (Donkey anti-Rabbit Secondary Antibody, Alexa Fluor 594, 1:1000, ThermoFisher) for one hour at room temperature, after which SFEBs were washed 3x with PBS with 0.1% Triton-X. SFEBs were incubated with 0.1 μg/ml Hoechst 33342 for 10 minutes at room temperature and washed once with PBS with 0.05% sodium azide. Prior to imaging, the plates were centrifuged at 163xg for 3 minutes and excess PBS was removed to facilitate auto-focusing.

ArrayScan XTI High Content Analysis Reader and the HCS Studio Cell Analysis Software (ThermoFisher Scientific) was used for high-content imaging analysis. This system is an automated image acquisition platform designed for confocal fluorescence imaging in microplates and is able to perform quantitative morphological measurements on a cell-by-cell or field-by-field basis. Due to the challenges of imaging SFEBs in suspension, and to account for variable thickness and size of SFEBs, microplate form factors were calibrated using average Z-focus obtained from several wells. Autofocusing was performed for each well using Hoechst 33342, with relaxed parameters and extended range of focus with 4×4 binning. Once autofocus was optimized, exposure times were calculated for each channel by setting the target percentile between 8-10%. Z-stacks were obtained for each channel in order to capture the entire range of the SFEB, and maximum intensity Z-projection was performed.

To analyze images, the neurite detection protocol was used on HCS studio. This protocol involves image processing; identification and validation of nuclei, cell bodies, and neurites; and analysis of neurite length and branching. During image processing, background is removed using a “low pass” filter to better resolve low-fluorescing bodies. For this study, nucleus identification was performed using Hoechst 33342 and analysis of cell bodies and neurites was performed using VGlut as a marker for excitatory neurons. Smoothing and dynamic thresholding was used to demarcate cell bodies and neurites in the SFEB. These two methods were used to overcome the differences in antibody penetration throughout the SFEB, which resulted in antibody intensity variability throughout the thickness of the SFEB. Other parameters such as the neurite identification modifier, direction length, point resolution, aggressive tracing and gap tolerance were all optimized to resolve cell bodies or neurites with weak fluorescent signals, dense fiber networks, and parallel neuronal fibers.

### Electrophysiology

In order to measure neuronal activity, Cytoview 48-well MEA plates (Axion Biosystems) were coated with 0.2% polyetylenimine (PEI) in borate buffer for 1 hour at room temperature, and washed 3x with distilled water. The plates were air-dried overnight, then a 10 μl droplet of laminin (20 μg/ml in terminal differentiation medium) was added directly on top of the recorded electrodes and incubated for an hour at 37°C. SFEBs were detached from cell culture inserts and transferred directly over the laminin droplet using a wide bore pipette. SFEBs were allowed to recover for one week prior to recording, with the media changed every other day.

Maestro MM and AxIS Software (Axion Biosytems) were used to measure spike frequency, bursting, and synchrony. Temperature was maintained at 37°C, and high and low pass filters were set to 200 Hz and 2000 Hz, respectively. The adaptive spike detection threshold was set to 5.25x standard deviation. Single electrode burst parameters were set to a minimum of 5 spikes per burst and a maximum inter-spike interval (Max ISI) of 100 ms. The synchrony window was set to 20 ms. Pharmacology was performed by recording from SFEBs before and after drug administration, and by comparing drug administration between control and disease lines. When necessary, drug washout was performed with two complete media changes. Advanced metrics were obtained via the Neural Metric Tool (Axion Biosystems). Experiments were repeated in triplicate and the following metrics were compared: mean firing rate (Hz), burst frequency (Hz), network burst frequency (Hz), and synchrony index. For the synchrony index, values closer to 1 represent higher synchrony. Continuous traces were generated through MATLAB.

To perform immunocytochemistry, SFEBs were fixed with 4% PFA for 45 minutes at room temperature. Staining was performed as mentioned above. SFEBs were imaged using the ArrayScan XTI at 10x magnification.

### Ca^2+^ imaging

SFEBs were transferred to 48-well MEA plates, transduced with pAAV9-CAG-GCamp6s-WPRE-SV40 (MOI=2.5 × 10^5^ Penn Vector Core; Addgene 100844-AAV9), and incubated for 6 days followed by a complete media change. Imaging was performed at least 7 days after viral transduction using the ThermoFisher Array Scan (HCI) at 5x magnification with a sampling of approximately 0.8 Hz. Analysis of calcium transients was performed using the Target Activation bioapplication on the HCS studio software suite. Average fluorescence intensity was recorded per region of interest (ROI) for every timepoint. The number and frequency of calcium signaling events were determined using PeakCaller^15^ (required rise/fall = 4%; max lookback/lookahead = 25 pts; trend control: finite difference diffusion (2-sided), trend speed = 60). PeakCaller was also used to generate raster plots.

### Statistics

Statistical analyses were performed using SigmaPlot 13.0 (Systat Software, Inc.). Normality was tested for all data-sets using the Shapiro-Wilk Test. For high-content imaging data, control and patient data were compared using the Mann-Whitney Rank Test. For electrophysiology, activity before and after drug administration was tested using the paired Wilcoxon-Signed Rank Test. All graphs were generated using Prism 6.0 (GraphPad Software Inc.)

## Results and Discussion

The SFEBs used in our system were generated from either control or patient cohorts. Upon maturing, SFEBs can be transferred to either a 48-well MEA plate allowing MEA recordings, single cell Ca^2+^ imaging, and immunocytochemistry, or a 96-well glass bottom plate for morphological studies. These SFEBs have telencephalon/forebrain characteristics and active glutamate/GABA synapses^10^. Neuronal markers showed that cells grow in an ‘inside out’ motif, similar to the development of the cortical subventricular zone^1^. High-throughput experimental techniques increase statistical power in this system and allow for detection and quantification of small differences between groups.

## Neuronal morphology

In order to quantify morphological changes between control and disease SFEBs, we performed high content screening using the ArrayScan XTI in SFEBs stained with the excitatory neuron marker VGlut with Hoechst as a nuclear stain. This method allows the detection of changes even in the presence of variability within each line. After scanning 96-well plates, images can be assayed at the level of individual wells, microscope field, or individual cells (Figure 2a,b). Masking protocols were used to measure multiple properties of VGlut positive (VGlut+) cells in control vs. disease cohorts. Total nuclei within the SFEB were decreased by 68%, (Figure 2c) in disease when compared to control, indicating that there were fewer cells per SFEB. When limiting measurements to excitatory neurons (VGlut+), the decrease was limited to 27% (Figure 2d). Additionally, the surface area of VGlut+ cells was also decreased by 50% (Figure 2e). In contrast, the disease model showed a 40% increase in VGlut+ neurite length (Figure 2d), with no significant difference in the number of branch points (Figure 2f). We observed a high degree of variability within SFEBs from the same line, and thus statistically significant changes may not be apparent using a lower sample size. Additionally, the method allows for analysis of several morphometric characteristics in a time- and cost-effective manner.

**Figure 2.**
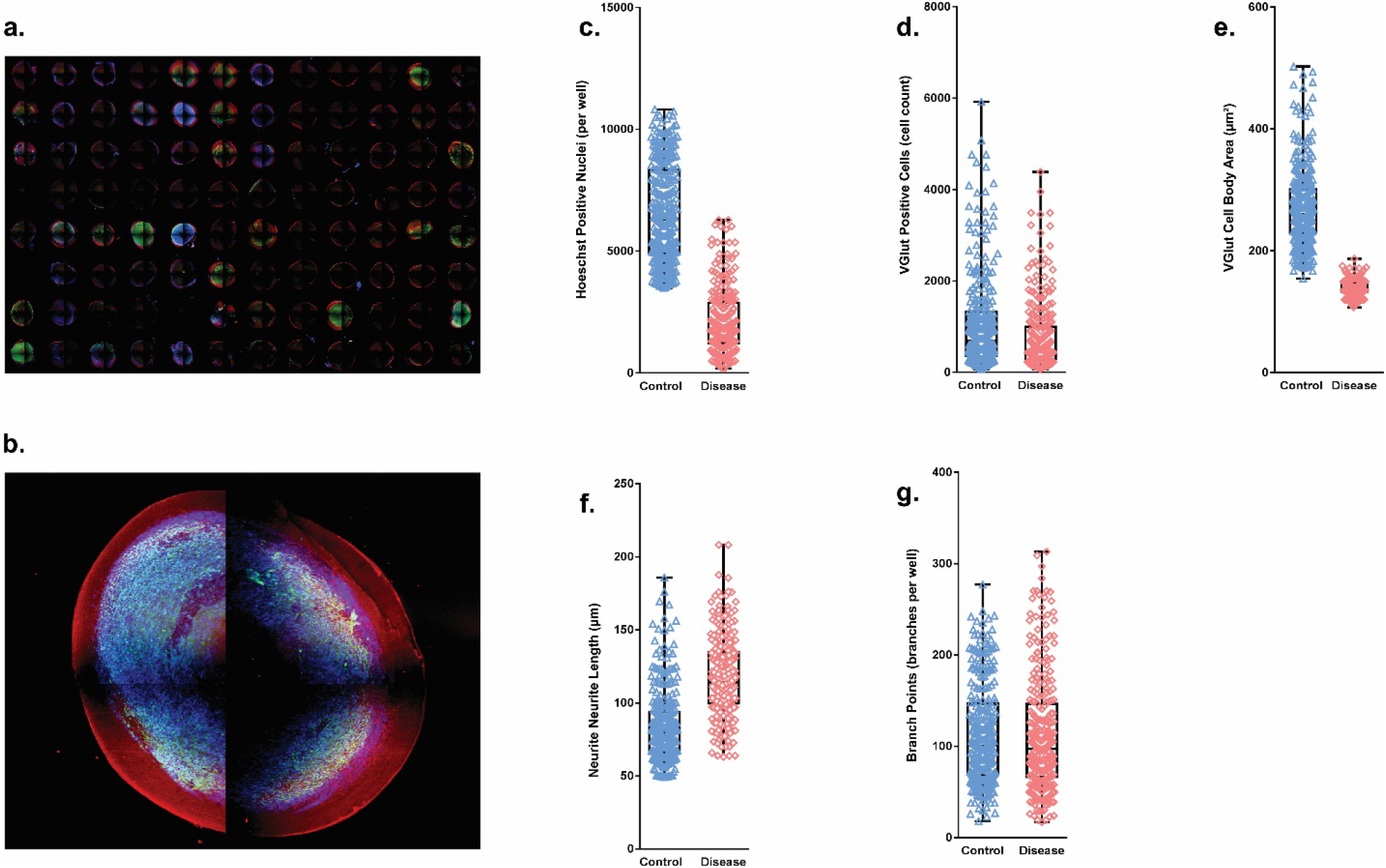
SFEB morphology measurements through ArrayScan screening platform. Following 7 days of recovery on optically clear 96-well U-bottom plates, SFEBs were stained for vesicular glutamate. (A) 96-well montage of SFEBs imaged using ArrayScan XTI. (B) High magnification image of SFEB. (C) Total number of Hoechst stained per SFEB was decreased by 68% in disease line (control = 6599.20 cells ±119.73 S.E.M; disease = 2134.961 cells ± 83.378 S.E.M.; p≤ 0.001; n = 285 control, 257 disease SFEBs, 3 expts; Mann-Whitney U). (D) Total number of excitatory neurons stained with VGlut was decreased by 26% (control = 1068.652 cells ± 63.765 S.E.M; disease = 780.498 cells ± 48.522 S.E.M.; p ≤ 0.001; n = 280 control, 262 disease SFEBs, 3 expts; Mann-Whitney U). (E) Mean cell body area of VGlut+ cells was decreased in disease line by 49% (control = 271.562 μm^2^ ± 3.974 S.E.M; disease = 139.744 μm^2^ ± 0.776 S.E.M.; p ≤ 0.001; n = 280 control, 277 disease SFEBs, 3 expts; Mann-Whitney U). (F) Total neurite length was increased by 43% in disease SFEBs (control = 84.388 μm ± 1.504 S.E.M; disease = 118.019 μm ± 1.662 S.E.M.; p ≤ 0.001; n = 286 control, 271 disease SFEBs, 3 expts; Mann-Whitney U). (G) Number of branch points in VGlut+ neurons is not significantly different between control and disease (control = 112.79 ± 3.244 S.E.M; disease = 112.609 ± 3.898 S.E.M.; p = 0.459; n = 286 control, 271 disease SFEBs, 3 expts; Mann-Whitney U).

A major limitation of imaging 3D cultures is that high cellular density and high refractive index of components within the cell membrane results in the scattering of light^16,17^. In the past, this has led to high-throughput screening techniques resulting in incomplete data sets and false negative data points^18^. Light scattering typically restricts imaging to the outer layers and restricts the use of more advanced physiological techniques such as microphotolysis and optogenetics. This method of growing the SFEBs in PTFE cell-culture inserts causes significant thinning compared to other 3D culture models^1^, allowing for greater penetration of light and limiting refraction. This allows for adoption of high content imaging techniques without the need for more labor-intensive processing such as sectioning, which is currently required in studying most 3D models^19^. Chemical clearing reagents can also be introduced in conjunction with this method to improve imaging by eliminating artifacts and high background fluorescence, although the effectiveness would need to be tested for individual markers. Tissue clearing protocols can also aid antibody penetration, which in turn allows for increased resolution of SFEB morphology.

## Network Firing Rates

SFEB local field potentials can be measured by MEA plates for multiple different endpoints. The most obvious are inherent differences in spontaneous activity between different groups, but bursting and network activity also can be examined. These measurements can be taken in the context of pharmacological manipulation of SFEB excitatory and inhibitory connections or modulatory neurotransmitter receptors. In this study, we used the NMDA antagonist APV (50μM) and the AMPA antagonist CNQX (25μM) to inhibit glutamatergic transmission. Both control and disease SFEBs show decreased mean firing rate and burst frequency in response to glutamatergic inhibition (Figure 3a). Application of the GABA antagonist picrotoxin (PTX, 50μM) increased the mean firing rate and burst frequency of control and disease SFEBs (Figure 3b). This shows significant excitatory and inhibitory neurotransmission in this culture model, consistent with previous findings indicating the presence of significant populations of excitatory and inhibitory neurons^10,11^. Synaptic plasticity, particularly long-term potentiation (LTP), also can be tested using this method. Addition of glycine has been shown to induce NMDA-dependent chemical LTP in cultured neurons^20^. Upon addition of glycine, we observed a significant increase in firing rate and burst frequency, which might be consistent with LTP induction (Figure 3c).

**Figure 3.**
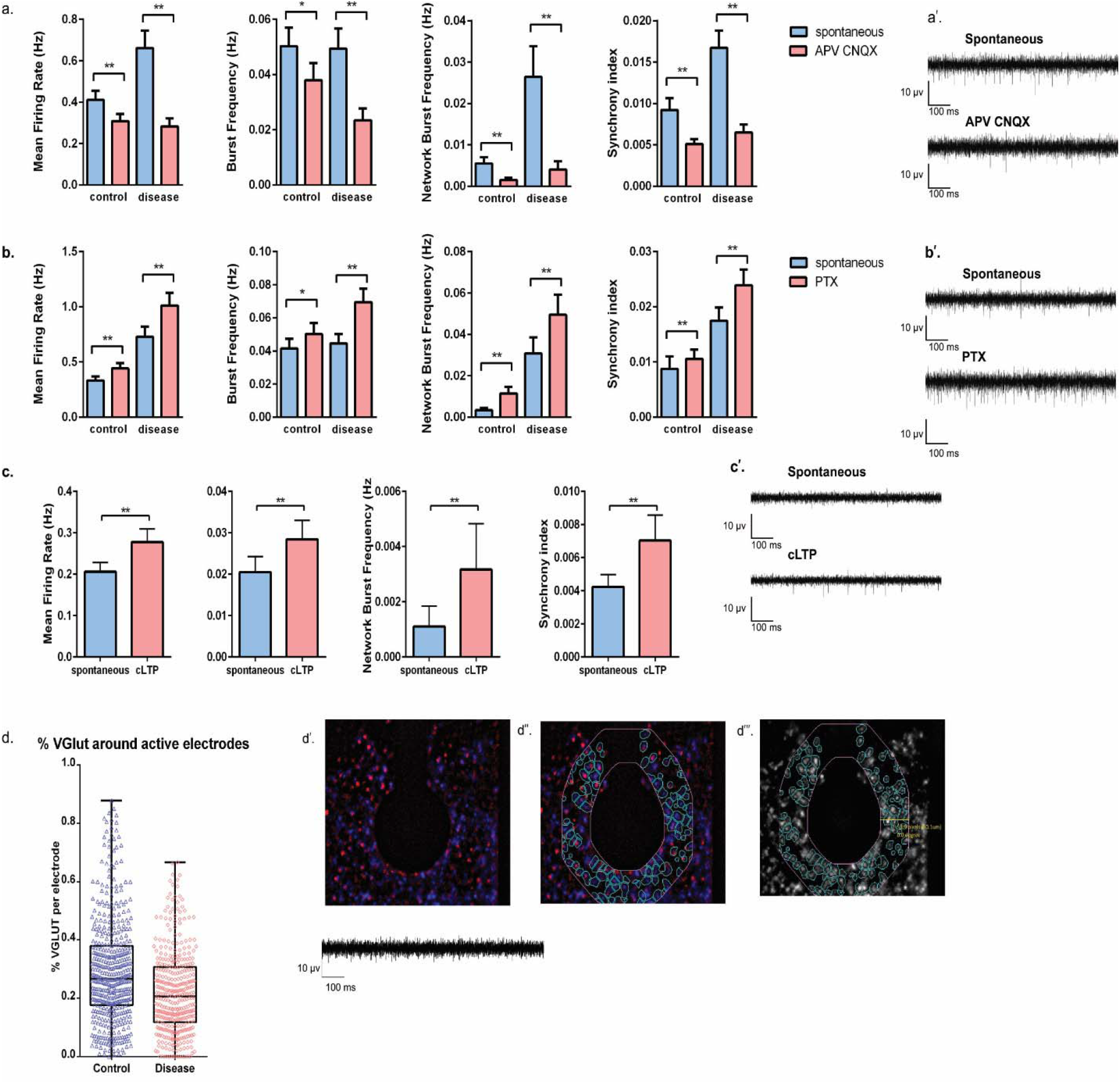
MEA analysis of SFEBs at DIV60 revealed significant differences between control and disease lines. (A) NMDA and AMPA inhibition with 50 μM APV and 25 μM CNQX decreased the mean firing rate of control SFEBs by 25% (spontaneous = 0.411 Hz ± 0.0448; APV CNQX = 0.309 Hz ± 0.0339; p < 0.001) and decreased the mean firing rate of disease SFEBs by 57% (spontaneous = 0.653 Hz ± 0.0829; APV CNQX = 0.279 Hz ± 0.038; p < 0.001). There was a 24% and 57% decrease in burst frequency in control and disease SFEBs, respectively (Control: spontaneous = 0.0503 Hz ± 0.00668; APV CNQX = 0.038Hz ± 0.00612; p = 0.020; Disease: spontaneous = 0.0494 Hz ± 0.00725; APV CNQX = 0.0234Hz ± 0.00425; p < 0.001). Network correlation as measured by frequency of network bursts and synchrony index. Network burst frequency was decreased by 72% in control lines (spontaneous = 0.00553 Hz ± 0.00153; APV CNQX = 0.00153 Hz ± 0.000489; p = 0.004) and 85% in disease lines (spontaneous = 0.653 Hz ± 0.0829; APV CNQX = 0.279 Hz ± 0.038; p < 0.001). Synchrony index was decreased by 45% in control (spontaneous = 0.00921± 0.00143; APV CNQX = 0.00511 ± 0.000578; p < 0.001) and 61% in disease SFEBs (spontaneous = 0.0167± 0.00209; APV CNQX = 0.00651± 0.000951; p < 0.001). All experiments were done in triplicate (n = 68 control, 71 disease SFEBs) and statistical analysis was done via the paired Wilcoxon Signed Rank Test. (B) Application of the GABA antagonist picrotoxin (PTX, 50 μM) increased the mean firing rate of control cells by 33% (spontaneous = 0.332 Hz ± 0.0364; PTX = 0.728 Hz ± 0.093; p < 0.001) with a 39% increase in the disease model (spontaneous = 0.441 Hz ± 0.0499; PTX = 1.012 Hz ± 0.113; p < 0.001). PTX also increased burst frequency in control SFEBs by 21% (spontaneous = 0.0415 Hz ± 0.00587; PTX = 0.0502 Hz ± 0.00679; p = 0.028) and in disease SFEBs by 56% (spontaneous = 0.0446 Hz ± 0.00564; PTX = 0.0695 Hz ± 0.00817; p < 0.001). Network burst frequency was increased by 2.3-fold in control (spontaneous = 0.00337 Hz± 0.00117; PTX = 0.0114 Hz ± 0.00315; p < 0.001) and 61% in disease SFEBs (spontaneous = 0.0308 Hz ± 0.00783; PTX = 0.0495 Hz ± 0.0096; p < 0.001). Synchrony was increased by 21% in control (spontaneous = 0.00875 ± 0.00224; PTX = 0.0106 ± 0.00168; p < 0.001) and 37% in disease SFEBs (spontaneous = 0.0175 ± 0.00237; PTX = 0.0239 ± 0.00283; p < 0.001). All experiments were done in triplicate (n = 68 control, 71 disease SFEBs) and statistical analysis was done via the paired Wilcoxon Signed Rank Test. (C) Chemical LTP was induced in SFEBs by application of 200 μM glycine. Treatment with glycine increased the mean firing rate control by 34% (spontaneous = 0.206 Hz ± 0.0224; cLTP = 0.277 Hz ± 0.0318; p < 0.001), burst frequency by 39% (spontaneous = 0.0205 Hz ± 0.00377; cLTP = 0.0506 Hz ± 0.00452; p = 0.005), network burst frequency by 2.9-fold (spontaneous = 0.00110 Hz ± 0.000738; cLTP = 0.00317 Hz ± 0.00166; p < 0.001), and synchrony index by 66% (spontaneous = 0.00423 ± 0.000744; cLTP = 0.00703 ± 0.00152; p < 0.001). All experiments were done in triplicate (n=128) and statistical analysis was done via the paired Wilcoxon Signed Rank Test. (D) In order to identify active electrodes, MEA recordings were performed and a threshold was set at a minimum of 5 spikes/min. Electrodes that met these criteria were imaged to assay the number of excitatory neurons that stained positive for VGlut within a 100 μm radius. This was done to examine the local microcircuitry surrounding active electrodes. The percentage of VGlut+ cells around active electrodes was lower in disease SFEBs (control = 0.2945 % ± 0.008272, n= 475 electrodes; disease = 0.2208 % ± 0.007208, n = 391 electrodes, p < 0.0001, Mann Whitney U).

Neuronal activity and local field potentials are largely dictated by local microcircuitry. To this end, we utilized our high content screening platform to analyze excitatory neuron populations surrounding individual electrodes. Using masking protocols, we were able to delineate an area of ~100 μm surrounding electrodes on a 48 well-MEA and quantify the percentage of VGlut+ cells surrounding active electrodes (at least 5 spikes per minute). Interestingly, although exhibiting higher spontaneous firing, disease SFEBs have less VGlut+ cells surrounding active electrodes compared to controls (Figure 3d). This may suggest altered inhibitory neuron populations or activity, or may indicate changes in the maturation state of excitatory neurons, which could point to interesting mechanisms underlying a disease phenotype.

## Calcium Signaling

Field recordings obtained by MEA can be supplemented by single-cell calcium imaging, which can provide information about the activity and location of individual neurons. For this method, we utilized the genetically encoded fluorescent calcium sensor GCaMP6s^13^ to visualize neural activity. Calcium transients were visible in SFEBs 7 days after induction with virus. High content imaging and analysis were performed using the ArrayScan XTI which allowed masking of individual cells and recording of average fluorescence over multiple time points (Figure 4A). To confirm that this activity originated from individual cells, individual electrodes were imaged at 20x. At this magnification, individual cells were clearly observed (Figure 4C). Analysis of high content fluorescence data using PeakCaller yielded the total number of fluorescent peaks, as well as the correlation of neuronal activity over time.

**Figure 4.**
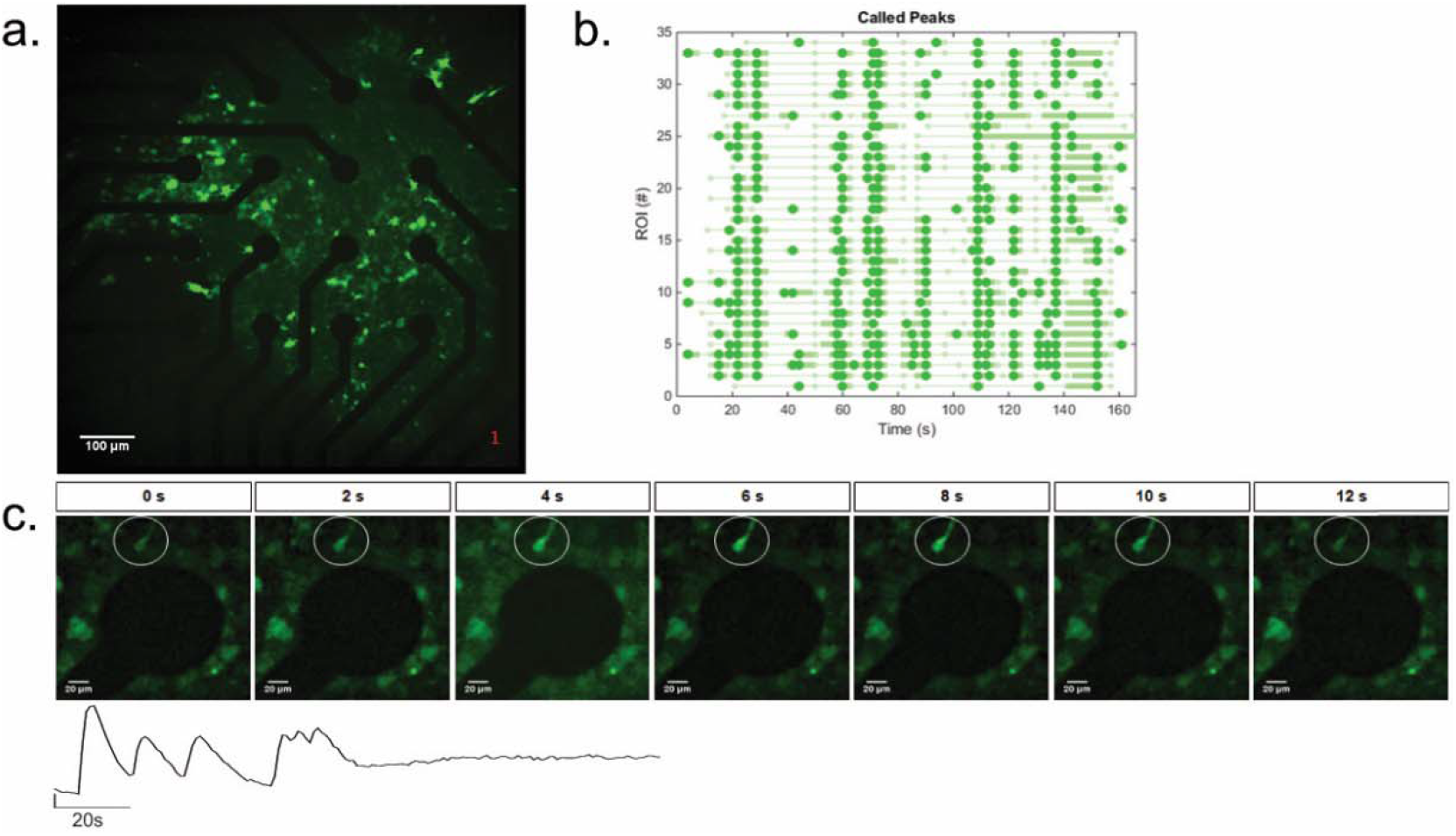
(A) High-throughput calcium imaging of SFEBs in 48-well MEA plates was performed using Arrayscan XTI at 5x magnification. (B) Raster plots were generated using PeakCaller from fluorescence intensity measurements obtained from an MEA well. (C) Activity in individual neurons were confirmed by imaging at higher magnification. Changes in fluorescence intensity can be clearly visualized (white circle).

Alternatively, single-cell activity can be studied via spike sorting, allowing spikes from putative neurons to be extracted from field recordings. This allows for a detailed examination of changes in spike properties from individual neurons such as amplitude, width, and spike-timing dependent plasticity.

## Conclusion

This workflow can be used to integrate high-throughput multiplex assays, heterogeneous patient population samples, and 3D culture to generate a vast amount of morphological and electrophysiological data that is more likely to reflect *in vivo* physiology^19^ compared to 2D cultures. SFEBs mimic early cortical development, and these assays may lead to a more complete understanding of the etiology of neurological disorders that originate developmentally. Additionally, the use of hiPSCs allows for the investigation of patient-specific pathophysiology. This approach can also be utilized for identification of potential drug targets and incorporated into a drug discovery pipeline. 3D cultures are more likely to reflect *in vivo* physiology compared to 2D cultures based on differences in cell proliferation, viability, gene expression profiles, and drug responses, and can potentially reduce development time, costs, and failure rates in drug discovery^19^. Ultimately, this is contingent upon further validation and standardization of techniques, and increased automation which can reduce experiment variability and improve cost-effectiveness. Overall, this method demonstrates the ability to gather rich and consistent quantitative data sets to detect differences between control and disease lines despite high variability inherent in 3D systems.

## Acknowledgements

This work is supported by a Maryland Stem Cell Research Fund Exploratory Grant MSCRFD-3815 and a Hussman Foundation Grant HIAS18002 to M.W.N. We thank Drs. Blatt and Hussman for helpful comments and review, as well as Elizabeth Benevides for review and Oscar Daniel Lara Montaño for the diagram in Figure 1.

## References

(1) Nestor, M. W.; Paull, D.; Jacob, S.; Sproul, A. A.; Alsaffar, A.; Campos, B. A.; Noggle, S. A. Differentiation of Serum-Free Embryoid Bodies from Human Induced Pluripotent Stem Cells into Networks. Stem Cell Res. 2013, 10 (3), 454–463. https://doi.org/10.1016/j.scr.2013.02.001.

(2) Duval, K.; Grover, H.; Han, L.-H.; Mou, Y.; Pegoraro, A. F.; Fredberg, J.; Chen, Z. Modeling Physiological Events in 2D vs. 3D Cell Culture. Physiology (Bethesda). 2017, 32 (4), 266–277. https://doi.org/10.1152/physiol.00036.2016.

(3) Clevers, H. Modeling Development and Disease with Organoids. Cell 2016, 165 (7), 1586–1597. https://doi.org/10.1016/j.cell.2016.05.082.

(4) Langhans, S. A. Three-Dimensional in Vitro Cell Culture Models in Drug Discovery and Drug Repositioning. Front. Pharmacol. 2018, 9 (JAN), 1–14. https://doi.org/10.3389/fphar.2018.00006.

(5) Wang, H. Modeling Neurological Diseases With Human Brain Organoids. Front. Synaptic Neurosci. 2018, 10 (June), 1–14. https://doi.org/10.3389/fnsyn.2018.00015.

(6) Tekin, H.; Simmons, S.; Cummings, B.; Gao, L.; Adiconis, X.; Hession, C. C.; Ghoshal, A.; Dionne, D.; Choudhury, S. R.; Yesilyurt, V.; et al. Effects of 3D Culturing Conditions on the Transcriptomic Profile of Stem-Cell-Derived Neurons. Nat. Biomed. Eng. 2018, 1– 15. https://doi.org/10.1038/s41551-018-0219-9.

(7) Pasca, A. M.; Sloan, S. A.; Clarke, L. E.; Tian, Y.; Makinson, C. D.; Huber, N.; Kim, C. H.; Park, J. Y.; O’Rourke, N. A.; Nguyen, K. D.; et al. Functional Cortical Neurons and Astrocytes from Human Pluripotent Stem Cells in 3D Culture. Nat. Methods 2015, 12 (7), 671–678. https://doi.org/10.1038/nmeth.3415.

(8) Carragher, N.; Piccinini, F.; Tesei, A.; Trask Jr, O. J.; Bickle, M.; Horvath, P. Concerns, Challenges and Promises of High-Content Analysis of 3D Cellular Models. Nat. Rev. Drug Discov. 2018, 17 (8), 606.

(9) Booij, T. H.; Price, L. S.; Danen, E. H. J. 3D Cell-Based Assays for Drug Screens: Challenges in Imaging, Image Analysis, and High-Content Analysis. SLAS Discov. Adv. Life Sci. R&D 2019, 247255521983008. https://doi.org/10.1177/2472555219830087.

(10) Phillips, A. W.; Nestor, J. E.; Nestor, M. W. Developing HiPSC Derived Serum Free Embryoid Bodies for the Interrogation of 3-D Stem Cell Cultures Using Physiologically Relevant Assays. J. Vis. Exp. 2017, No. 125. https://doi.org/10.3791/55799.

(11) Nestor, M. W.; Jacob, S.; Sun, B.; Prè, D.; Sproul, A. A.; Hong, S. I.; Woodard, C.; Zimmer, M.; Chinchalongporn, V.; Arancio, O.; et al. Characterization of a Subpopulation of Developing Cortical Interneurons from Human IPSCs within Serum-Free Embryoid Bodies. Am. J. Physiol. Cell Physiol. 2015, 308 (3), C209–19. https://doi.org/10.1152/ajpcell.00263.2014.

(12) Brennand, K. J.; Marchetto, M. C.; Benvenisty, N.; Brüstle, O.; Ebert, A.; Izpisua Belmonte, J. C.; Kaykas, A.; Lancaster, M. A.; Livesey, F. J.; McConnell, M. J.; et al. Creating Patient-Specific Neural Cells for the in Vitro Study of Brain Disorders. Stem Cell Reports 2015, 5 (6), 933–945. https://doi.org/10.1016/j.stemcr.2015.10.011.

(13) Chen, T. W.; Wardill, T. J.; Sun, Y.; Pulver, S. R.; Renninger, S. L.; Baohan, A.; Schreiter, E. R.; Kerr, R. A.; Orger, M. B.; Jayaraman, V.; et al. Ultrasensitive Fluorescent Proteins for Imaging Neuronal Activity. Nature 2013, 499 (7458), 295–300. https://doi.org/10.1038/nature12354.

(14) DeRosa, B. A.; Van Baaren, J. M.; Dubey, G. K.; Lee, J. M.; Cuccaro, M. L.; Vance, J. M.; Pericak-Vance, M. A.; Dykxhoorn, D. M. Derivation of Autism Spectrum Disorder-Specific Induced Pluripotent Stem Cells from Peripheral Blood Mononuclear Cells. Neurosci. Lett. 2012, 516 (1), 9–14. https://doi.org/10.1016/j.neulet.2012.02.086.

(15) Artimovich, E.; Jackson, R. K.; Kilander, M. B. C.; Lin, Y. C.; Nestor, M. W. PeakCaller: An Automated Graphical Interface for the Quantification of Intracellular Calcium Obtained by High-Content Screening. BMC Neurosci. 2017, 18 (1), 1–15. https://doi.org/10.1186/s12868-017-0391-y.

(16) Hell, S. W.; Reiner, G.; Cremer, C.; Stelzer, E. H. K. Aberrations in Confocal Fluorescence Microscopy Induced by Mistakes in Refractive Index. J Microsc 1993, 169 (March), 391–405.

(17) Jacques, S. L. Optical Properties of Biological Tissues◻: A Review. Phys. Med. Biol. 2013, 37 (58), R37–R61. https://doi.org/10.1088/0031-9155/58/11/R37.

(18) Pampaloni, F.; Ansari, N.; Stelzer, E. H. K. High-Resolution Deep Imaging of Live Cellular Spheroids with Light-Sheet-Based Fluorescence Microscopy. Cell Tissue Res. 2013, 352 (1), 161–177. https://doi.org/10.1007/s00441-013-1589-7.

(19) Boutin, M. E.; Voss, T. C.; Titus, S. A.; Cruz-Gutierrez, K.; Michael, S.; Ferrer, M. A High-Throughput Imaging and Nuclear Segmentation Analysis Protocol for Cleared 3D Culture Models. Sci. Rep. 2018, 8 (1), 1–14. https://doi.org/10.1038/s41598-018-29169-0.

(20) Musleh, W.; Bi, X.; Tocco, G.; Yaghoubi, S.; Baudry, M. Glycine-Induced Long-Term Potentiation Is Associated with Structural and Functional Modifications of Alpha-Amino-3-Hydroxyl-5-Methyl-4-Isoxazolepropionic Acid Receptors. Proc. Natl. Acad. Sci. U. S. A. 1997, 94 (17), 9451–9456. https://doi.org/10.1073/pnas.94.17.9451.

